# Beyond Motility: A Multiparametric Fitness Assay Framework to Evaluate Anthelmintics in *Nippostrongylus brasiliensis*

**DOI:** 10.1101/2025.06.12.659374

**Authors:** Kyle Anesko, Sarah Midou, Rebecca Ruggiero-Ruff, Yuxin He, Meritte Lotfy, Jennell Jennett, Meera G. Nair, Adler R. Dillman

## Abstract

We adapted a multiparametric fitness assay panel—including ATP quantification, larval growth, pigmentation, and cuticle permeability—to assess *Nippostrongylus brasiliensis* viability *in vitro*. To validate this system, we benchmarked two known anthelmintics with distinct modes of action: quinidine, a heme detoxification disruptor, and pyrantel pamoate, a neuromuscular antagonist. We then applied this framework to NCR247, a plant-derived heme-sequestering peptide with proposed antiparasitic activity. While the assay panel sensitively detected physiological disruption by known anthelmintics, NCR247 did not significantly impair parasite fitness across any tested parameter. These findings highlight the utility of integrated viability assays and suggest further *in vivo* or delivery-optimized studies may be necessary to evaluate NCR247 efficacy.

## Introduction

Hookworm infections remain one of the most significant causes of neglected tropical disease (NTD), affecting over 700 million people worldwide, particularly in impoverished communities with limited access to sanitation and healthcare [1]. Chronic infection can lead to iron-deficiency, anemia, malnutrition, and developmental impairments [2,3]. Despite their global impact, effective and lasting interventions for hookworm remain elusive. Albendazole—the most commonly used anthelmintic—shows variable efficacy and increasing signs of resistance [4–7]. Although vaccine candidates are under investigation, none have yet reached clinical application [9–11].

A major barrier to advancing hookworm research is the limited availability of *Necator americanus* for experimental studies [8]. Consequently, *Nippostrongylus brasiliensis* (Nb), a rodent parasite with a similar life cycle, tissue tropism, and blood-feeding behavior, has become a valuable laboratory model for preclinical studies [8,9]. Notably, Nb shares conserved pathways with human hookworms, including those involved in heme acquisition and detoxification, making it a promising surrogate for drug screening [9].

Traditional *in vitro* hookworm assays primarily rely on parasite motility as a proxy for viability. However, these assessments are often subjective, low-throughput, and do not reflect physiological status [10]. To overcome these limitations, we developed a complementary panel of viability assays that quantify ATP levels, larval growth, intestinal pigmentation, and cuticle permeability—four metrics that collectively provide a more nuanced view of parasite fitness.

To validate this assay panel, we benchmarked its sensitivity using two known anthelmintics with distinct modes of action: pyrantel pamoate, a neuromuscular disruptor, and quinidine, a heme-detoxification inhibitor. We then applied this framework to evaluate the D-enantiomer of NCR247, a heme-sequestering plant peptide originally identified in legume symbiosis, where it sequesters heme to manipulate *Sinorhizobium meliloti*’s iron regulatory pathways—not by depriving the bacterium of essential heme, but by causing it to perceive iron starvation and upregulate iron import [11–13]. Because heme is achiral, the D-enantiomer sequesters heme identically to the naturally occurring L-enantiomer but is protease resistant [11]. Given the heme auxotrophy of hookworms, we hypothesized that NCR247 would impair parasite viability by limiting access to this essential nutrient. This study thus serves both to evaluate the efficacy of NCR247 and to demonstrate the broader utility of multiparametric fitness assays in antiparasitic screening.

## Materials and Methods

### Ethics statement

All procedures involving animals were approved by the University of California, Riverside institutional animal care and use committee (IACUC) (protocols A-20150028B and A-20150027E) and were in accordance with National Institutes of Health guidelines, the animal welfare act, and public health service policy on humane care and use of laboratory animals.

### Animal husbandry

Sprague-Dawley rats raised in a microbiologically defined environment (SD-F MPF) female rats were purchased from Taconic Biosciences and maintained in-house. Rats were age-matched for Nb-infections (6-8 weeks old) and housed two per cage under an ambient temperature with a 12 h light/12 h dark cycle. When infected with Nb iL3s, rats were monitored for clinical signs of infection or distress on day 2, 3, and 5 post-infection. C57BL/6 mice purchased from the Jackson Laboratory were bred in-house. Mice were age matched (6 to 14 weeks old), gender matched and housed five per cage under an ambient temperature with a 12 h light/12 h dark cycle.

### *Nippostrongylus brasiliensis* (Nb) Collection and Preparation

*Nippostrongylus brasiliensis* (Nb) iL3s were obtained from rats. About 5000 infective third-stage larvae (iL3s) of *N. brasiliensis* were subcutaneously injected into SD-F MPF rats to induce Nb infection and continue their life cycle. Fecal cultures were prepared 6–10 days post-infection using activated charcoal and vermiculite in petri dishes. Plates were aerated daily and lightly moistened three times per week with sterile water. All Nb iL3s used in experiments were age matched (2-3 weeks old), and isolated using a modified Baermann technique.[14] Isolated iL3s were washed in sterile 1x phosphate-buffered saline (1× PBS), then exsheathed by adding 5 volumes of bleach solution (4.8 mL PBS and 200 µL 5.65 - 6% sodium hypochlorite solution (Fisher Scientific)) for up to 1 minute while vortexing, then washed twice with sterile 1× PBS, followed by a 1-hour incubation in antibiotic solution (0.012 g neomycin and 1.2 mL penicillin/streptomycin [100×] brought to 30 mL with sterile 1× PBS). Worms were counted under a dissecting microscope and then maintained at room temperature in sterile DMEM supplemented with 10% Heat-inactivated FBS, 1% penicillin/streptomycin, 1mM Sodium Pyruvate, and 25 mM HEPES at room temperature. Nb iL3s were maintained in supplemented DMEM at room temperature for no longer than 6 hours prior to experimentation.

### Red Blood Cell Collection

Red blood cells (RBCs) were isolated from C57BL/6 mice via cardiac puncture or mesenteric artery under terminal isoflurane anesthesia, and anticoagulated in Alsever’s solution (2.05% dextrose (Sigma), 0.8% sodium citrate (Sigma), 0.055% citric acid (Sigma), and 0.42% sodium chloride (Sigma) in double distilled H_2_O). After collection, whole blood was washed three times in RBC Wash Buffer (21 mM tris(hydroxymethyl)aminomethane (Sigma), 4.7 mM KCl (Sigma), 2.0 mM CaCl_2_ (Sigma), 140.5 mM NaCl (Sigma), 1.2 mM MgSO_4_ (Sigma), 5.5 mM glucose (Sigma), and 0.5% bovine albumin fraction V (BSA)), with the serum being removed after each wash to isolate RBCs.[9] Then RBCs were resuspended in sterile 1× PBS and counted using a hemocytometer. RBCs were maintained in 1× PBS at 4°C for up to 2 days before use in experiments.

### *In Vitro* Culture Conditions

Unless otherwise stated, all treatment groups were co-cultured with 1×10^8^ mouse red blood cells (RBCs) per well in 48-well plates, incubated in supplemented DMEM at 37 °C with 5% CO_2_ for 4 days.

### *Nb* Morphology & Pigmentation

iL3s were incubated with 10^8^ RBCs per well in 48-well plates at 37 °C with 5% CO_2_ for up to 7 days. Worms were imaged using a Keyence microscope to assess morphology and pigmentation. Worm morphology was assessed by tracing length and width in brightfield images using ImageJ. Worm length was determined by tracing the worm lengthwise along the central line. Worm width was measured by tracing the widest region of the midsection. All measurements were normalized to a control group of iL3s incubated in supplemented DMEM without RBCs or drugs. Pigmentation change was calculated by normalizing each group’s Day 4 darkness to its respective Day 0 baseline. Relative darkness was measured by converting images to 16-bit grayscale in ImageJ and using ImageJ’s freehand tool to trace the worm’s perimeter and measure mean gray value. Each biological group consisted of 3 technical replicates, with 15–30 Nb iL3s imaged per replicate and averaged.

### ATP Assay

On day 4 post-incubation, residual RBCs were lysed with sterile water to remove debris before collecting iL3s. After collection, iL3s were washed three times with 1× PBS, counted and aliquoted into tubes at 100 iL3s per 150 µL 1× PBS with a single 6.35 mm steel bead. They were homogenized using a bead beater at 4°C for 5 min. The homogenates were centrifuged at 1000 × *g* for 2 min. Supernatants were transferred to a black 96-well clear-bottom plate and mixed with 1 volume of CellTiter-Glo 2.0 reagent (Promega) per well. Luminescence was measured using a SpectraMax i3 plate reader. ATP values were calculated using a standard curve prepared with ATP disodium salt, and ATP levels were normalized to worm count to yield per-worm ATP values. Each biological group consisted of 3 technical replicates, with 100 Nb iL3s being measured per replicate and averaged.

### Cuticle Permeability

On day 4 post-incubation, media was replaced with 1× PBS. iL3s were stained with 100 µM SYTOX Green and incubated for 24 hours at 37 °C in 5% CO_2_. After staining, worms were washed and imaged on a Keyence BZ-X series fluorescence microscope. Cuticle permeability was quantified using ImageJ by tracing each worm’s perimeter with the freehand selection tool and calculating the mean gray value. Each biological replicate consisted of three technical replicates, with approximately 30 Nb iL3s imaged per replicate and averaged.

### Statistical Analysis

All experiments were conducted in biological triplicate, with each replicate representing an independent experimental run performed on a separate day. For each run, treatment groups were analyzed using one-way ANOVA with Dunnett’s post hoc test relative to matched internal controls. No statistical comparisons were made across experiments with differing treatment compositions. Data are presented as mean ± SEM unless otherwise stated. Statistical significance was defined as p < 0.05. Analyses were performed using GraphPad Prism v10.4.

## Results

### Benchmarking and evaluation of compound effects on ATP levels in Nippostrongylus brasiliensis

To validate the sensitivity and specificity of our multi-assay platform, we first tested two known anthelmintics—quinidine (QN) and pyrantel pamoate (Pp)—before evaluating the effects of the D-enantiomer of NCR247 on ATP levels in *Nippostrongylus brasiliensis* [9].

We next applied the assay to NCR247, here referring to the D-enantiomer form, a heme-sequestering peptide hypothesized to impair parasite viability by limiting access to host-derived heme. In a dose-response experiment, NCR247 was added to RBC-fed cultures at concentrations ranging from 0.1 µM to 2.5 mM. Across the lower range (0.1–100 µM), no significant reduction in ATP levels was observed compared to vehicle-treated controls (Fig. 1B). However, at concentrations ≥1 mM, ATP levels were significantly reduced. Light microscopy revealed that at these concentrations, NCR247 formed a visible gel-like matrix in the media, physically restricting parasite movement and likely reducing viability through nonspecific entrapment rather than a true biological mechanism—especially since ATP reduction was also observed under RBC-free conditions. To avoid this artifact, concentrations ≥1 mM were excluded from further in vitro testing. Among the usable doses, 10 µM NCR247 yielded the lowest ATP values, and was therefore selected as the working concentration for subsequent experiments.

**Figure 1.**
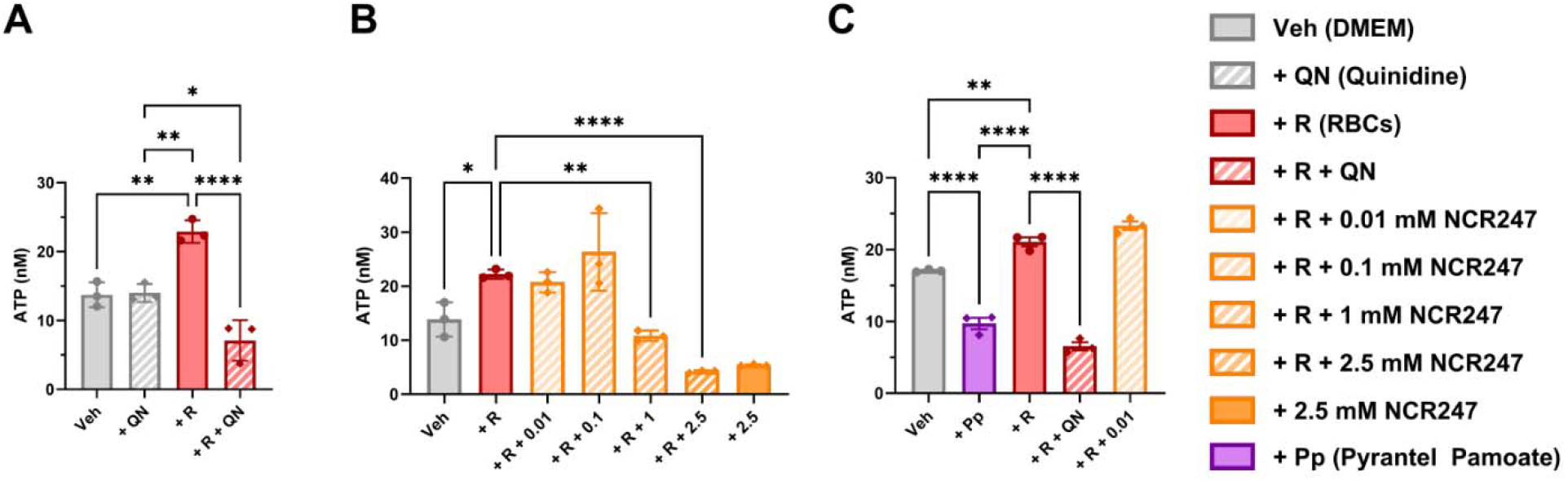
ATP-based assessment of blood-dependent and blood-independent anthelmintic activity in *N. brasiliensis*. All experiments assessed ATP levels in approximately 100 N. brasiliensis iL3s after 4 days of in vitro culture under the indicated treatment conditions. ATP levels were measured using CellTiter-Glo 2.0. **(A)** Assay validation with Quinidine (QN, 10µM), a heme-detoxification disruptor. QN reduced ATP levels only in larvae cultured with red blood cells (+R, 1×10^8^ RBCs), confirming blood-feeding–dependent activity. **(B)** Dose-response curve of the D-enantiomer of NCR247in RBC-fed larvae. ATP levels were reduced only at concentrations ≥1 mM, which were excluded from interpretation due to media gelation and physical entrapment. **(C)** Validation with Pyrantel pamoate (Pp, 10µM), a neuromuscular anthelmintic. Pp significantly reduced ATP levels regardless of RBC presence, confirming blood-independent activity.Data are presented as mean ± SEM. Statistical analysis was performed using ordinary one-way ANOVA with Tukey’s multiple comparison test. Only statistically significant comparisons are shown; all others were not significant (ns). Data represent three biological replicates, each with three technical replicates per group.

In contrast, 10 µM Pp and 10 µM QN significantly reduced ATP levels through distinct mechanisms: Pp in an RBC-independent manner and QN in an RBC-dependent manner (Fig. 1C), confirming the assay’s sensitivity to multiple anthelmintic modes of action. ATP levels were used as a proxy for parasite viability and metabolic fitness. All compounds were delivered in supplemented DMEM, which served as the vehicle control in all groups.

### Morphology-based assay detects stunted larval growth from Pyrantel and Quinidine, but not the D-enantiomer of NCR247

To assess the sensitivity of our assay to developmental disruption, we evaluated changes in larval morphology following drug exposure. *Nippostrongylus brasiliensis* iL3s were cultured in vitro for 4 days with the indicated treatments, and worm length and width were measured using brightfield microscopy. RBC-fed controls exhibited a modest increase in both dimensions, consistent with normal growth and molting during early development (Fig. 2A–B). Known anthelmintics with cytotoxic or neuromuscular effects, including Pyrantel pamoate (Pp) and Quinidine (QN), significantly reduced larval size, confirming the assay’s ability to detect impaired growth. In contrast, the D-enantiomer of NCR247(10 µM) had no significant effect on either parameter compared to RBC-fed controls, suggesting it does not disrupt larval development under these conditions.

**Figure 2.**
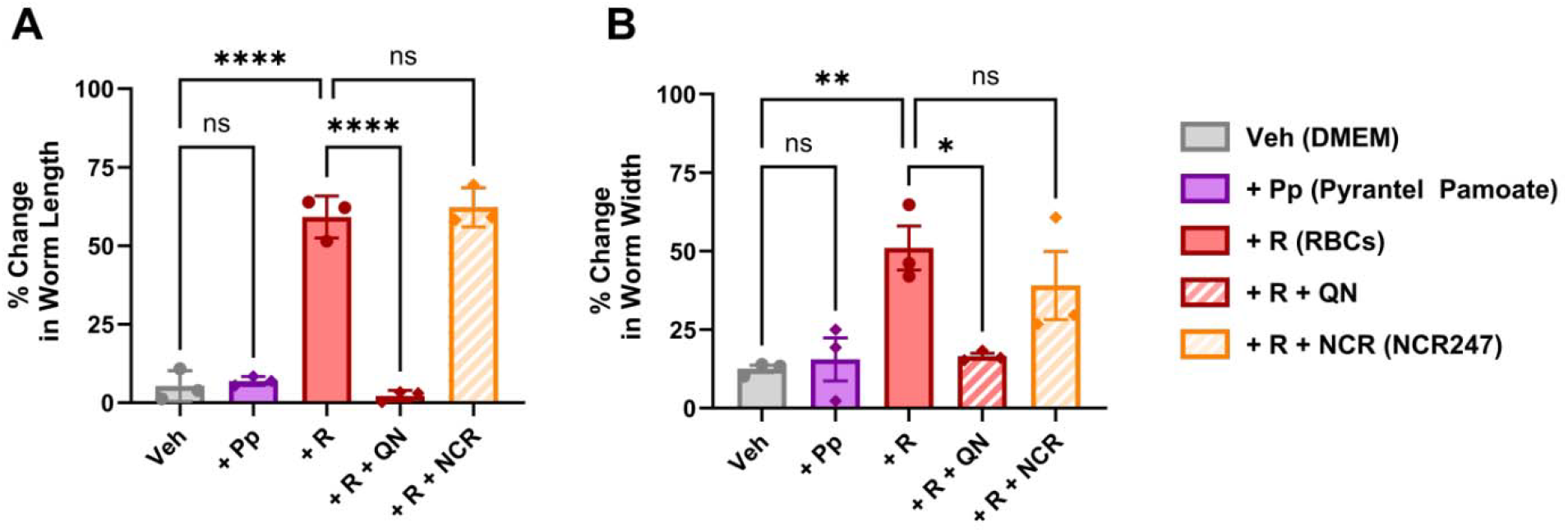
Comparative evaluation of anthelmintics shows selective inhibition of *N. brasiliensis* growth. All experiments measured percent change in length and width of Nippostrongylus brasiliensis iL3s after 4 days of in vitro culture with the indicated treatments. **(A, B)** Percent change in worm length and width, respectively, was quantified using brightfield micrographs analyzed in ImageJ. Red blood cells (+R, 1 × 10^8^ RBCs) supported normal larval growth. Pyrantel pamoate (+Pp, 10 µM) and Quinidine (+QN, 10 µM) significantly reduced larval dimensions. D-enantiomer of NCR247(+R + NCR, 10 µM) had no measurable effect on growth. Data are shown as mean ± SEM from three biological replicates, each containing three technical replicates per group. Statistical analysis was performed using one-way ANOVA with Tukey’s multiple comparisons test. Only statistically significant differences are indicated; all others were not significant (ns).

To determine the optimal exposure period for morphological analysis, we performed a time-course experiment from Day 0 to Day 7. Measurable increases in worm length and width were observed by Day 4, with minimal changes beyond that point (Supplemental Fig. 1). These data support the use of a 4-day exposure period as an appropriate endpoint for detecting growth-modifying effects.

ATP-based assays were conducted in parallel to evaluate metabolic effects (Fig. 1).

Together, these findings demonstrate that Pp and QN impair both parasite energy metabolism and developmental progression, while NCR247 (D-enantiomer) does not affect either under physiologically relevant conditions.

### Quinidine and Pyrantel impair pigmentation in *N. brasiliensis*, but the D-enantiomer of NCR247 does not disrupt blood ingestion

To evaluate whether the D-enantiomer of NCR247 affects blood ingestion or heme accumulation, we quantified pigmentation in *Nippostrongylus brasiliensis* iL3s after 4 days of co-culture with red blood cells (RBCs). As positive controls, we included quinidine (QN), which interferes with heme detoxification, and pyrantel pamoate (Pp), which disrupts feeding behavior. Both QN and Pp significantly reduced pigmentation, validating the assay’s sensitivity to disruptions in heme utilization and ingestion (Fig. 3C). In contrast, the D-enantiomer of NCR247(10 µM) did not significantly affect pigmentation relative to RBC-fed controls, suggesting it does not interfere with blood ingestion or downstream heme handling under these conditions.

**Figure 3.**
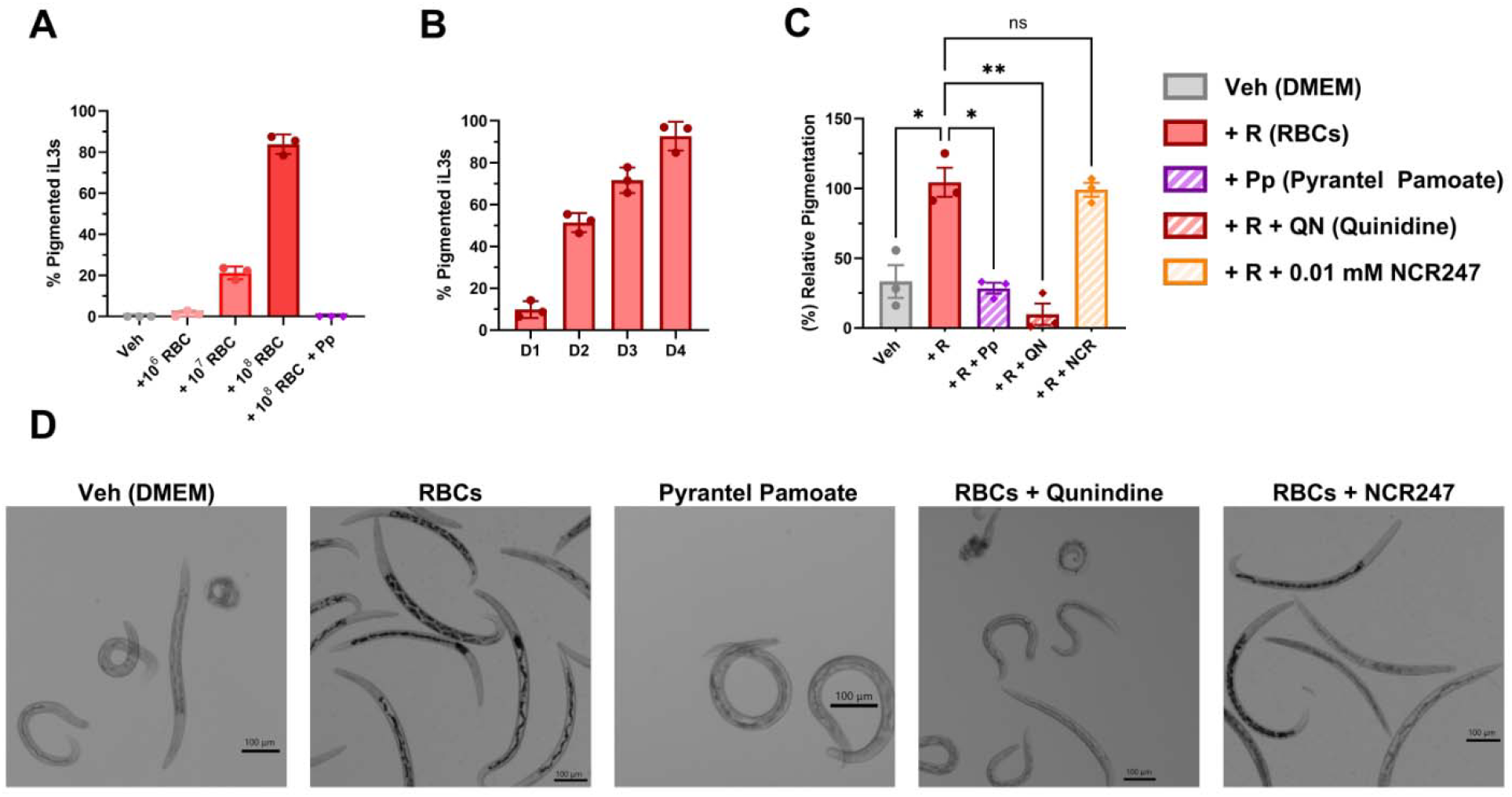
Pigmentation analysis reveals impaired heme utilization from benchmark anthelmintics, but not the D-enantiomer of NCR247. All experiments assessed pigmentation levels in *N. brasiliensis* iL3s after 4 days of in vitro culture under the indicated treatment conditions. Pigmentation was measured using brightfield microscopy and quantified as relative image darkness (grayscale intensity) in ImageJ. **(A)** RBC dose-response curve: Worms were cultured with increasing RBC concentrations (10^6^–10^8^ RBCs) for 4 days. **(B)** Time-course of pigmentation in worms exposed to 10L RBCs for 1–4 days. **(C)** Pigmentation in worms co-cultured with RBCs (10^8^) and treated with Quinidine (QN, 10LµM), Pyrantel pamoate (Pp, 10µM), or the D-enantiomer of NCR247(10 µM). Pigmentation values were normalized to each group’s Day 0 baseline to account for batch variability. **(D)** Representative brightfield images showing pigmentation patterns across groups. Scale bars = 100 µm. Data are presented as mean ± SEM (n = 100 worms per group; 3 biological replicates, 3 technical replicates per biological replicate). Statistical analysis was performed using Brown–Forsythe and Welch ANOVA to account for unequal variance. Statistically significant comparisons are shown; all others are not significant (ns).

To establish optimal assay parameters, we first characterized pigmentation intensity across a range of RBC concentrations and exposure durations. Pigmentation increased in both a dose-dependent (Fig. 3A) anda time-dependent manner (Fig. 3B), reaching saturation by Day 4 in worms cultured with 10^8^ RBCs. These findings confirmed that Day 4 and 10^8^ RBCs represent suitable benchmark conditions for assessing blood-feeding–dependent pigmentation.

### Cuticle permeability assay detects structural disruption from Pyrantel and Quinidine, but not the D-enantiomer of NCR247

To validate our cuticle permeability (CP) assay, we performed a dose-response analysis using Pyrantel pamoate (Pp), a compound previously shown to disrupt parasite structural integrity [15,16]. As expected, SYTOX Green uptake increased with rising concentrations of Pp, and worms cultured in boiled DMEM exhibited maximal fluorescence (Fig. 4A), confirming the assay’s ability to detect cuticular compromise.

**Figure 4.**
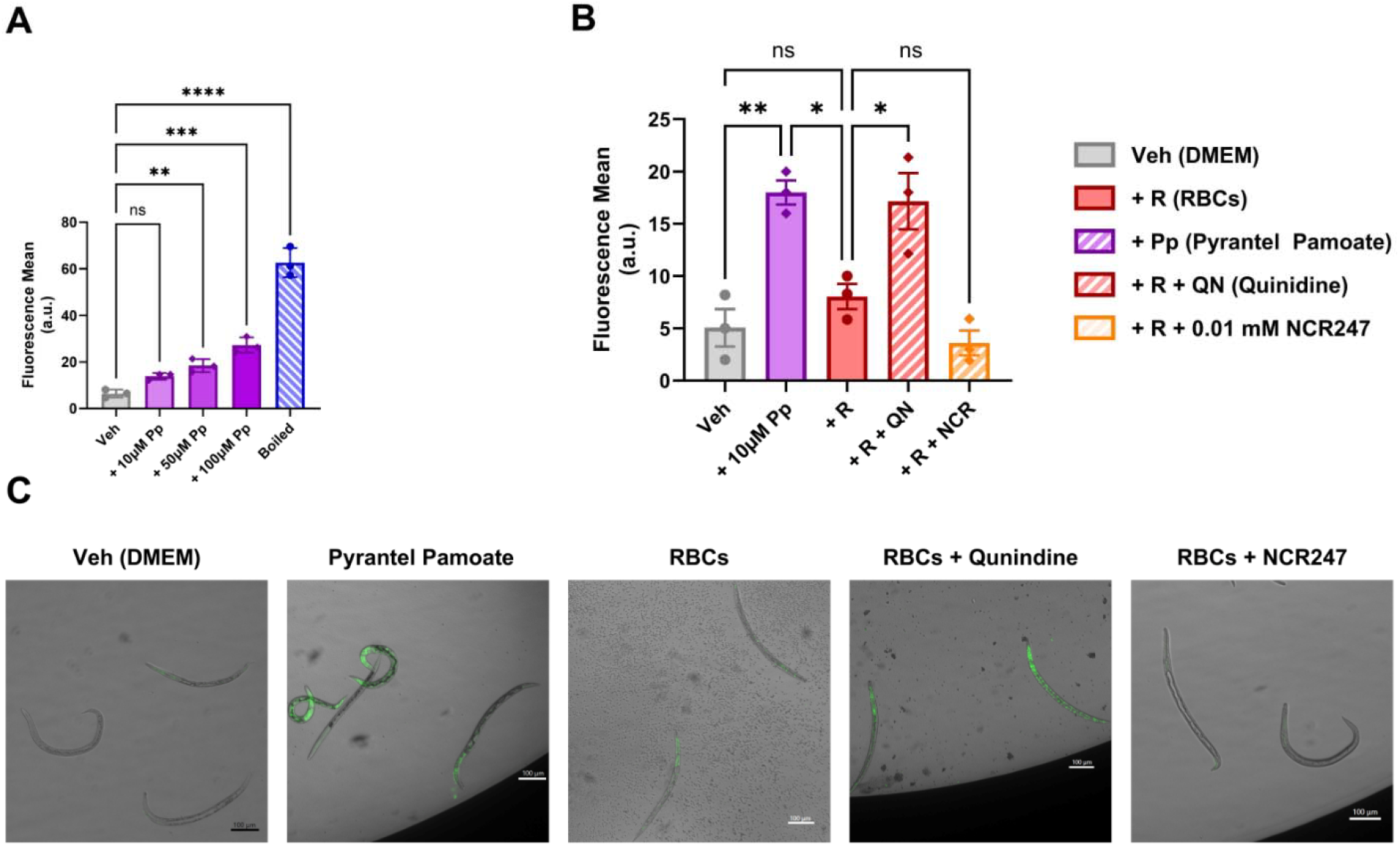
Pyrantel and Quinidine increase cuticle permeability in *N. brasiliensis*, but the D-enantiomer of NCR247 does not affect structural integrity. Cuticle permeability (CP) in *N. brasiliensis* iL3s was assessed after 4 days of in vitro culture using a SYTOX Green uptake assay. Fluorescence intensity per worm was quantified using ImageJ. **(A)** Dose-response analysis of Pyrantel pamoate (Pp) to validate assay performance. Worms were treated with increasing concentrations of Pp (1–100 µM), or cultured in boiled DMEM (10 minutes) as a positive control for surface damage. **(B)** CP in worms treated with Pyrantel pamoate (Pp), Quinidine (QN), or the D-enantiomer of NCR247(10 µM). All treatment groups were co-cultured with 10^8^ RBCs, except for the Pp-only condition. **(C)** Representative fluorescence images showing SYTOX Green uptake. Increased fluorescence reflects cuticle damage. Scale bars = 100 µm. Data are presented as mean ± SEM (n = 100 worms per group; 3 biological replicates, 3 technical replicates per replicate). Statistical analysis was performed using one-way ANOVA with Tukey’s multiple comparison test. Statistically significant comparisons are shown; all others are not significant (ns).

We then applied the assay to assess whether the D-enantiomer of NCR247 disrupts larval surface integrity. As benchmarking controls, Pp and Quinidine (QN) significantly increased SYTOX fluorescence, indicating sensitivity to both mechanical and heme-disruptive mechanisms (Fig. 4B). In contrast, D-enantiomer of NCR247-treated larvae exhibited fluorescence comparable to RBC-fed controls, suggesting that NCR247 does not compromise cuticle integrity under these in vitro conditions.

## Discussion

In this study, we developed and validated a multiparametric assay panel to assess the physiological fitness of *Nippostrongylus brasiliensis* iL3s in vitro. By integrating quantitative readouts for energy metabolism (ATP), morphology, pigmentation, and cuticle permeability, we captured a broad range of viability metrics beyond traditional motility scoring. This framework enabled sensitive detection of known anthelmintic effects and facilitated evaluation of the D-enantiomer of NCR247, hereby referred to as NCR247, a heme-sequestering peptide with potential antiparasitic activity. Benchmarking with quinidine and pyrantel pamoate confirmed the sensitivity and versatility of this platform. Pyrantel pamoate impaired ATP production and growth and increased cuticle permeability, consistent with its systemic toxicity. Quinidine selectively reduced ATP and pigmentation in RBC-fed worms, in line with a heme-detoxification–dependent mechanism. These findings affirm the ability of our assays to distinguish between compounds with distinct modes of action.

In contrast, NCR247 did not significantly alter any physiological fitness metric at concentrations of 10 µM or less, despite its known ability to bind heme with high affinity [11]. Several explanations may account for this apparent lack of activity. First, NCR247 acts by sequestering free heme extracellularly, and its efficacy depends on achieving sufficient stoichiometric excess to limit parasite access to this essential nutrient. Our assay conditions included co-culture with 1×10^8^ mouse RBCs per well (∼1% hematocrit), chosen to ensure robust feeding and pigmentation within a 4-day window. Even assuming ideal 1:1 binding, 10 µM NCR247 would be insufficient to significantly deplete this heme pool. Thus, the peptide’s sequestration capacity was likely outcompeted by the high heme burden present in vitro.

Notably, although our hematocrit was lower than physiological blood levels (typically 35–45%), the absolute number of RBCs—and thus the total heme content—remained extremely high, potentially exceeding the heme concentrations encountered by parasites in the host gut [9].

Second, NCR247’s extracellular mechanism may be inherently less disruptive than that of intracellular-acting compounds. Quinidine, for example, is membrane-permeable and likely disrupts intracellular heme detoxification, leading to the accumulation of toxic free heme [11,17,18]. In contrast, NCR247 does not damage the parasite directly but instead limits heme availability in the environment. Because sequestered heme is not inherently toxic, parasites may remain viable if their internal heme stores or uptake pathways are sufficient [19–22].

Furthermore, *N. brasiliensis* may possess compensatory mechanisms—such as heme storage, redundant import pathways, or stage-specific adaptations—that buffer against transient heme limitation, particularly in the iL3 stage.

Finally, experimental limitations may also contribute. Although the D-enantiomer is protease-resistant, peptide instability due to oxidation or nonspecific binding to serum proteins or plastic surfaces cannot be excluded [11]. The 4-day culture period may also be insufficient to reveal delayed viability effects resulting from progressive heme deprivation. Additionally, we found that at concentrations ≥1 mM, NCR247 aggregated into a gel-like matrix, precluding experimentation at those concentrations. Future work should consider testing NCR247 under reduced heme conditions, in later developmental stages with increased heme demand, or in combination with permeabilizing agents that enhance peptide retention or cellular stress. There may also be modifications or structural variants of NCR247 that exhibit reduced aggregation at higher concentrations, which would facilitate additional experimentation.

Despite these findings, this study highlights the value of a multiparametric fitness framework in parasitology research. Compared to traditional motility assays, this platform enables higher-resolution phenotyping, supports mechanistic inference, and is adaptable for medium-throughput drug screening. Future applications could expand this approach to additional life stages, parasite species, or host–parasite co-culture systems. Coupling these assays with motility tracking or 3D invasion models may offer further insight into compound effects on parasite behavior and survival.

In summary, while the D-enantiomer of NCR247 did not significantly impair *N. brasiliensis* iL3 viability under the conditions tested, the multiparametric assay system developed here provides a powerful tool for anthelmintic discovery and functional screening. This approach may help identify new compounds and uncover vulnerabilities that are not detectable with single-metric assays.

## Acknowledgements

We thank Dr. Graham Walker and his laboratory at the Massachusetts Institute of Technology for generously providing the D-enantiomer of NCR247 peptide used in this study. We are also grateful to Dr. Walker for his insightful feedback and editorial suggestions.

---

**Supplemental Figure 1.**
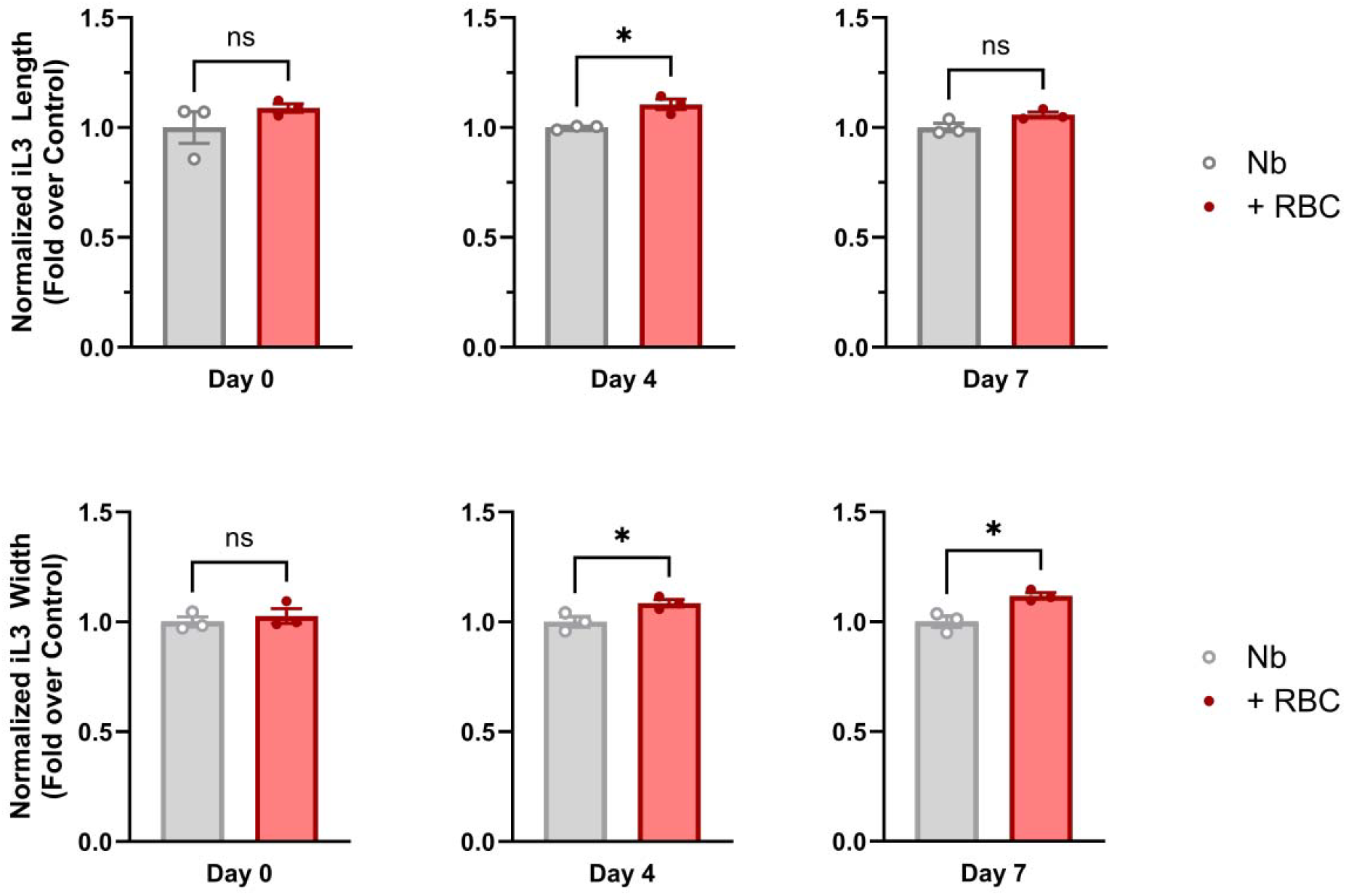
Blood feeding promotes *Nippostrongylus brasiliensis* growth and development *in vitro*. **(A)** Representative brightfield images of *N. brasiliensis* iL3 larvae cultured in vitro with or without red blood cells (RBCs) for up to 7 days. Images show larvae at Day 0 and Day 4. **(B)** Quantification of worm length at Day 0, Day 4, and Day 7. **(C)** Quantification of worm width at Day 0, Day 4, and Day 7. Worms were cultured in media alone (Nb) or media supplemented with 10^8^ RBCs (+ RBCs). Length and width were measured using ImageJ. Data are presented as mean ± SEM (n = 15 worms per group), with three biological replicates and three technical replicates per biological replicate. Statistically significant differences were observed between Nb and Nb + RBCs groups at both Day 4 and Day 7. Statistical analysis was performed using one-way ANOVA with Tukey’s post hoc test.

